# Insights into gadolinium uptake and release dynamics of a macrocyclic contrast agent in blood cells

**DOI:** 10.64898/2026.07.08.736994

**Authors:** Alberto Cornet-Gomez, Nathalie Peyer, Lena Sophia Zaugg, Liesel Goveas, Cristina Zivko, Peter Broekmann, Johannes T. Heverhagen, Hendrik von Tengg-Kobligk, Nico Ruprecht

## Abstract

**Background:** Gadolinium-based contrast agents (GBCAs) are routinely used in magnetic resonance imaging (MRI). Although macrocyclic GBCAs were initially considered biologically inert, it is now known that a fraction of patients retains gadolinium (Gd) for prolonged periods in tissues such as blood, bone, and brain. Because the first cellular interactions of GBCAs occur in the bloodstream, this study aimed to elucidate the uptake mechanism but also the intracellular persistence and release dynamics of gadoterate meglumine, one of the most widely used macrocyclic agents, in white blood cells (WBCs).

**Methodology and principal findings:** WBCs and K562 cells were incubated with gadoterate meglumine under different conditions to investigate its cellular entry mechanisms. Uptake of the contrast agent was quantified by measuring intracellular Gd using single-cell inductively coupled plasma mass spectrometry (SC-ICP-MS). Time and concentration-dependent incubation of K562 cells revealed saturable uptake kinetics consistent with a Michaelis–Menten model which is independent of the phase of the cell cycle. Gadoterate meglumine uptake in both WBCs and K562 cells was shown to be an active process, as uptake was strongly reduced or abolished at low temperature (16°C and 4°C) and in the presence of metabolic inhibitors (sodium azide and 2-deoxyglucose). Co-incubation with multiple endocytosis inhibitors (Dyngo 4a, Dynole 2-24 and chlorpromazine) did not significantly decrease intracellular Gd levels in K562 cells and caused only a slight reduction in WBCs, indicating that endocytosis is not the main entry pathway for gadoterate meglumine in these cells. Furthermore, we assessed the retention time of the Gd inside the cells, showing that only after 24 hours post incubation 80% percent of the intracellular Gd was released through an active process. Finally, we demonstrate that one of the mechanisms of Gd release from WBCs involves extracellular vesicles, which may substantially increase its potential for downstream accumulation in different tissues, including immunoprivileged tissues like brain.

**Significance:** The observed time-dependent accumulation, temperature and energy dependence of gadoterate meglumine uptake demonstrate that active cellular mechanisms are primarily responsible for GBCA internalization. Furthermore, our results indicate that macropinocytosis, phagocytosis, and clathrin-mediated endocytosis are not the primary routes of gadoterate meglumine entry. Hereby, we also describe that Gd externalization is an active process involving extracellular vesicles which may influence the Gd distribution in different tissues and its consequent long-term retention. Further studies are required to explore strategies to block this process in order to mitigate potential long-term gadolinium retention.

## Introduction

The theoretical principles and first human scans using magnetic resonance imaging (MRI) were developed in the 1970s, however, the MRI was formally introduced into the clinical arena in the early 1980s with the development and installation of the first commercial whole-body scanners (1). MRI is now widely adopted as both a clinical and research tool as a non-invasive imaging technique that generates cross-sectional images of internal structures using non-ionizing electromagnetic radiation (2). While powerful, MRI is fundamentally limited by its low signal-to-noise ratio. To overcome this limitation, gadolinium-based contrast agents (GBCAs) are frequently administered to improve the spatial resolution. Since the first GBCA received Food and Drug Administration (FDA) approval in 1988, over 750 million doses have been administered worldwide (3, 4). GBCAs consist of a highly toxic trivalent gadolinium ion (Gd^3+^) enclosed by an organic ligand. This chelation with an aminopolycarboxylic acid ligand is critical because the ion’s similarity to calcium allows it to interfere with key biological processes (5). The GBCAs used in the clinic are composed of Gd^3+^ conjugated to either linear or macrocyclic ligands. These complexes generally show high thermodynamic and kinetic stability constants (6, 7).

By the early 2000s, concerns about GBCAs began to grow. While the risk of acute allergic reactions was previously known, a more severe concern emerged in 2006 when GBCAs were first linked to the pathogenesis of nephrogenic systemic fibrosis (NSF) (8, 9). Key studies linked patients with acute renal failure who were given the linear GBCA Magnevist with an increased likelihood of developing NSF (9). Sanyal et al. documented the widespread tissue deposition of insoluble Gd-phosphate in a post-mortem analysis of an NSF patient, with deposits found in multiple organs including the skin, liver, lungs, and cerebellum thereby confirming the extensive nature of Gd accumulation in affected individuals (10). This finding aligns with a study showing that fibroblasts and macrophages have demonstrated that after endosomal uptake, the acidic lysosomal environment facilitates the degradation of some linear GBCAs leading to the deposition of Gd as phosphates within hours of cellular entrapment (11).

Growing evidence indicates that GBCAs can leave trace gadolinium-containing deposits in the brain and other tissues, in both humans and animal model studies with normal renal function (12–14). These deposits could be related with pathological symptoms, known as symptoms associated with gadolinium exposure (SAGE) or even gadolinium deposition disease (GDD) (15, 16). Despite the lack of a definitively established link between these deposits and adverse health effects, these findings have prompted a re-evaluation of the GBCA safety profile and resulted in revised guidelines from regulatory authorities on their use (17). The implementation of these new regulatory frameworks has not only led to a reduction in the incidence of NSF (14) but has also opened new avenues for assessing the potential long-term consequences of Gd accumulation, such as its effects on motor and cognitive functions (18).

Previous findings of our lab showed that WBCs from some patients and healthy donors exhibited heterogeneous intracellular Gd concentrations following GBCA administration (19). These observations led to the central hypothesis of the present study: immune cells may act as carriers of GBCAs, thereby facilitating their transport and contributing to gadolinium deposition in the body (20). To test this hypothesis, it is essential to understand how GBCAs are internalized and released from WBCs. The distinction between active and passive transport is crucial for a complete understanding of GBCA pharmacology. Passive transport, such as simple or facilitated diffusion, would suggest movement down a concentration gradient without direct energy dependence, potentially influenced by the GBCA’s physicochemical properties (21). In contrast, active transport which includes endocytosis (e.g., pinocytosis) or specific transporter-mediated uptake, implies an energy-dependent process capable of moving GBCAs against a concentration gradient (22). Identifying the dominant transport mechanism and the intracellular retention time will provide vital insights into the downstream biological consequences within immune cells and other tissues.

In this work we aimed to elucidate the mechanisms underlying the gadoterate meglumine uptake and release by WBCs. Gadoterate meglumine is one of the most widely used GBCAs in clinical practice; in this study, Dotarem (Guerbet, Villepinte, France) was used to perform the experiments. To overcome the high inter-individual variability observed in patients, we used K562 cells as a controlled experimental model, a human cell line derived from a patient with chronic myelogenous leukemia. Furthermore, the most relevant findings were validated in WBCs isolated from consented healthy donors.

## Results

### Uptake of gadoterate meglumine is time and concentration-dependent exhibiting saturable kinetics in K562 cells

To uncover if K562 are able to uptake gadoterate meglumine in a concentration and time-dependent manner. Fig 1A shows that K562 cells incubated with Gadoterate meglumine at concentrations ranging from 0.2-10 mM for 60 min at 36 °C, increased with extracellular GBCA concentration and approached a plateau at higher concentrations, demonstrating a saturable uptake profile. The obtained curve fits to the Michaelis-Menten equation yielding a V_max_ of 1.2×10^−11^ mol⋅g protein^−1^⋅s^−1^ and a K_m_ of 1.3 mM. This saturable profile suggests that Gd uptake in K562 cells is an active, transporter-mediated process, rather than simple passive diffusion.

**Figure 1.**
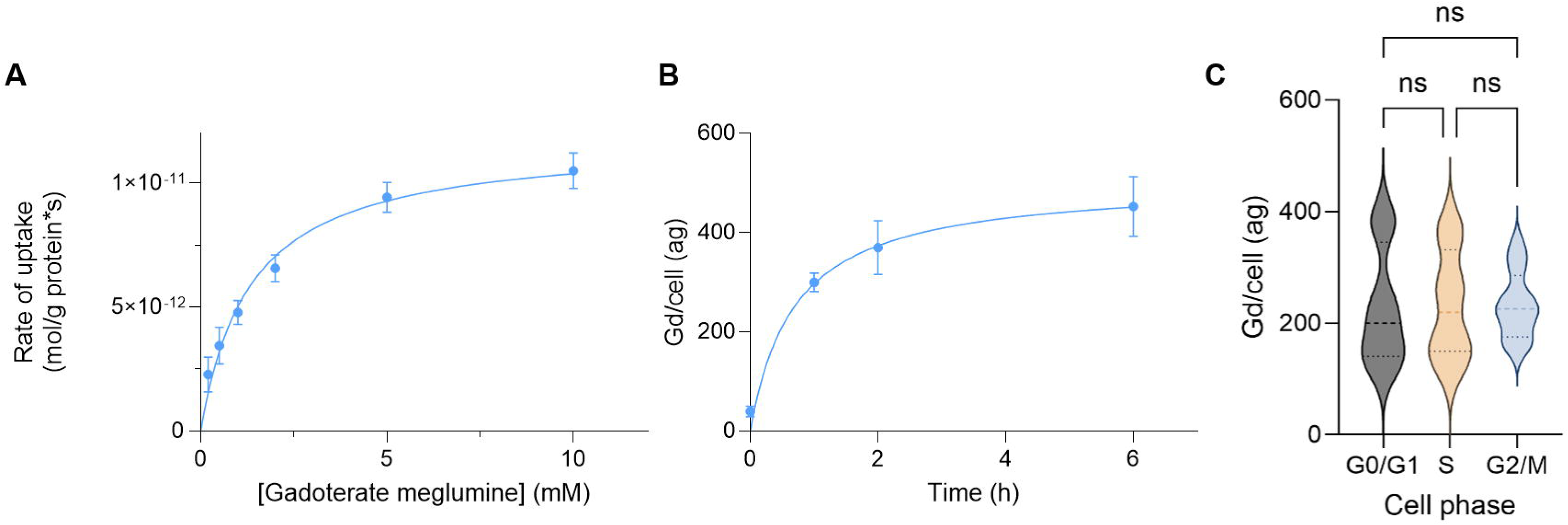
Michaelis-Menten kinetics of gadoterate meglumine uptake in K562 cells. (A) Concentration-dependent uptake of gadoterate meglumine in K562 cells. This figure illustrates a saturable uptake of process consistent with Michaelis–Menten kinetics. Each data point represents the mean of three independent experiments, with error bars indicating the standard deviation. Fitting the data to the Michaelis-Menten equation in GraphPad Prism yielded a Vmax of 1.2×10^−11^ mol⋅g protein^−1^⋅s^−1^ and a Km of 1.3 mM. (B) Time-dependent uptake of gadoterate meglumine in K562 cells. Uptake shows an initial linear increase between 0 and 2 h, followed by a plateau phase in which intracellular Gd levels no longer increase proportionally with time. Each data point represents the mean of three independent experiments, with error bars indicating the standard deviation. (C) Gadoterate meglumine uptake is independent on the cell phase in which the K562 cells are, they can internalize the molecule in every stage in a similar manner. Statistical analysis was performed using one-way ANOVA followed by Bonferroni’s post hoc test.

Based on our results (Fig. 1A) and considering that in the clinics, where patients are administered GBCAs intravenously at a concentration of 2 mM for contrast enhanced MRIs, we decided to maintain 2 mM of gadoterate meglumine as the working concentration for all the experiments. Then, Fig. 1B show the K562 uptake dependence with the time of incubation. Gadoterate meglumine uptake by these cells shows an initial linear increase between 0 and 2 h, followed by a plateau phase in which intracellular Gd levels no longer increase proportionally with time. Taking into account these data and the fact that most of the patients with normal renal function present an elimination half-life of less than 2 hours, the experiments in this work were performed with 1 hour incubation.

Furthermore, we aimed to determine whether uptake was dependent on the phase of the cell cycle in which cells were at the time of stimulation. In this regard, Figure 1C shows that K562 cells internalize gadoterate meglumine to a similar extent during the G0/G1, S, and G2/M phases, indicating that this process is independent of the cell cycle.

### Evidence for temperature dependent uptake of GBCAs in K562 cells and WBCs

Our investigation revealed a clear temperature-dependent uptake of gadoterate meglumine by both K562 cells (Fig. 2A) and WBCs (Fig. 2B). At the physiological temperature of 36 °C, both K562 cells and WBCs exhibited significant gadoterate meglumine uptake, whereas minimal intracellular Gd was detected at 4 °C. Receptor-mediated endocytosis and vesicular trafficking are known to be highly temperature dependent, with internalization being nearly abolished below 10-15 °C and markedly reduced at approximately 16-20 °C. Consistent with this, gadoterate meglumine uptake at 16 °C was intermediate between that observed at 36 °C and 4 °C. The similar values obtained upon both the 4°C and 16°C incubations are not consistent with a vesicular uptake. Together, the rate and extent of intracellular Gd accumulation quantified by SC-ICP-MS strongly indicate that Gd uptake is mediated by active cellular mechanisms rather than by passive diffusion.

**Figure 2.**
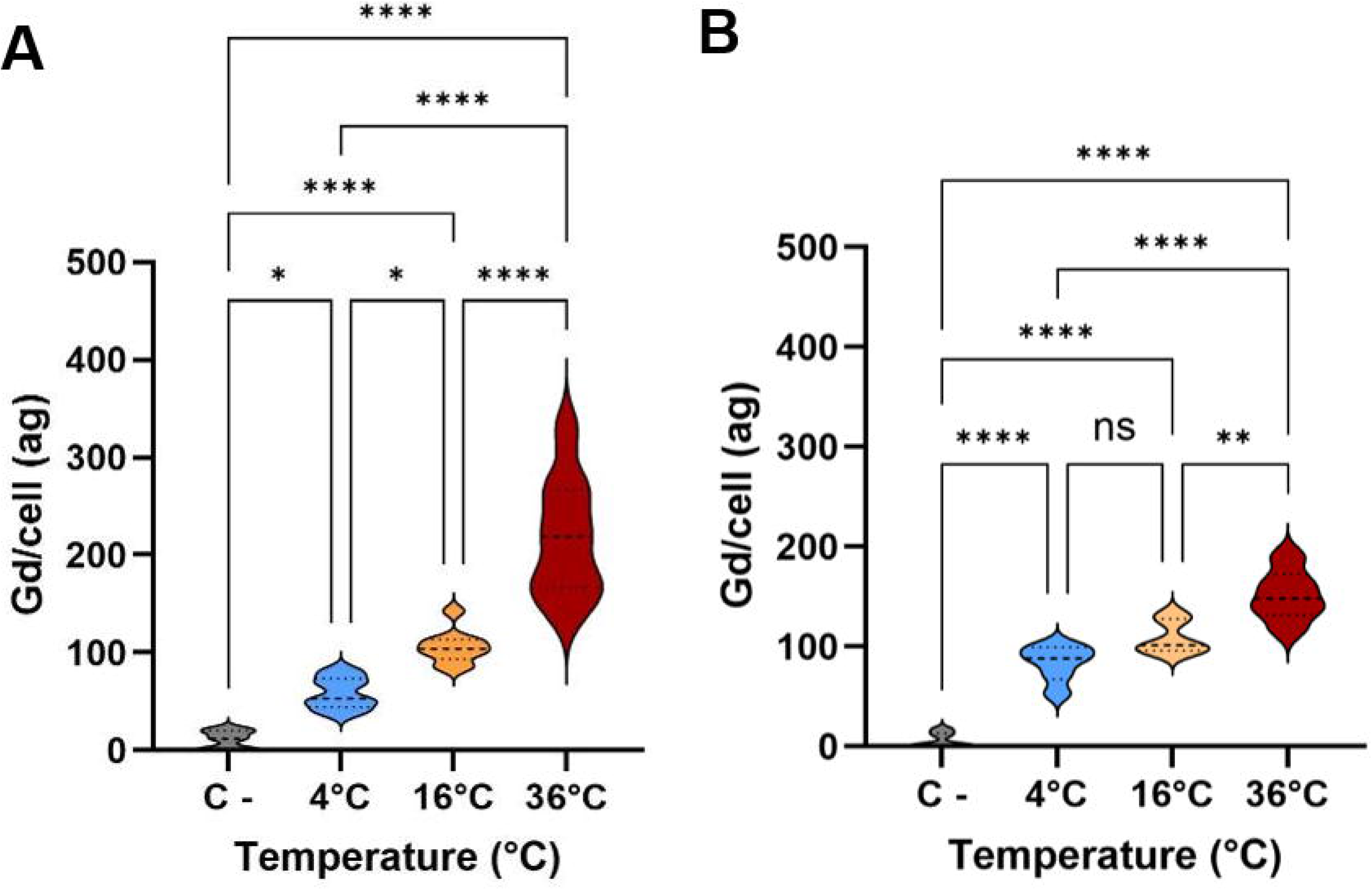
Temperature-dependent uptake of gadoterate meglumine in K562 cells (A) and WBCs (B). Cells were incubated with 2 mM gadoterate meglumine for 1 hour at 4 °C (blue), 16 °C (orange), or 36 °C (red). Gd uptake was temperature dependent, with maximal uptake at 36 °C, intermediate uptake at 16 °C, and minimal uptake at 4 °C. Cells not incubated with gadoterate meglumine were used as negative controls (grey). Data are shown as mean ± SD of three biological replicates. Statistical analysis was performed using one-way ANOVA followed by Bonferroni’s post hoc test. *P < 0.05, **P < 0.01, ****P < 0.0001.

### Complete metabolic inhibition reveals energy-dependent uptake in K562 cells and WBCs

To definitively confirm that GBCA uptake is mediated by active, energy-dependent mechanisms, we induced ATP depletion by inhibiting oxidative phosphorylation and glycolysis with sodium azide (NaN₃) and 2-deoxy-D-glucose (2DG), respectively (23, 24). Under these energy-depleted conditions, gadolinium uptake was significantly reduced in both K562 cells and WBCs (Fig. 3) compared to cells incubated with 2 mM gadoterate meglumine alone (C+), thereby reinforcing the conclusion that gadoterate meglumine uptake is an energy-dependent process in these cells.

**Figure 3.**
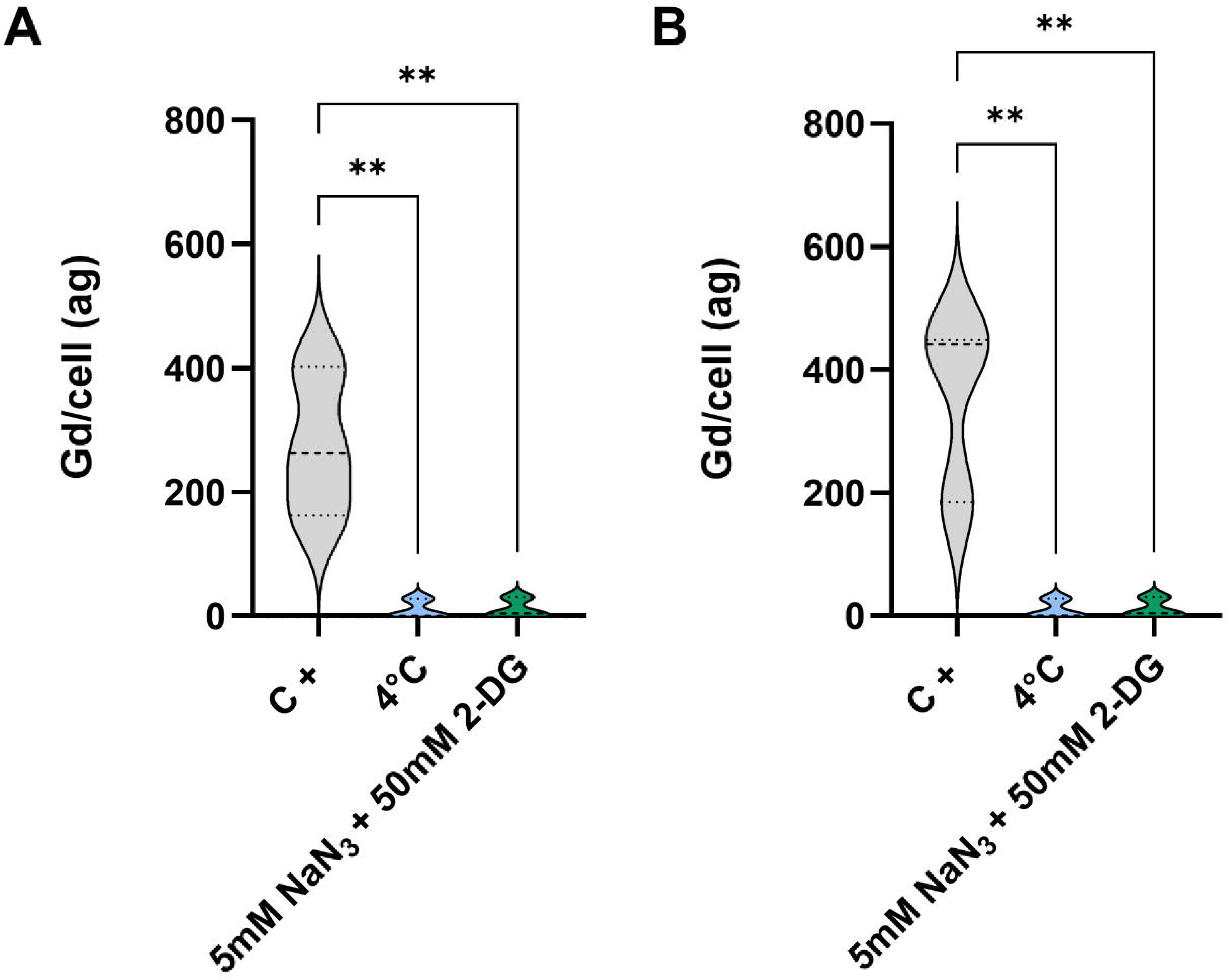
ATP depletion inhibits gadoterate meglumine uptake in K562 cells (A) and WBCs (B). Cells were treated with 5 mM NaN₃ and 50 mM 2-deoxy-D-glucose to deplete intracellular ATP (green). Under these conditions, gadoterate meglumine uptake was strongly reduced in both K562 cells and WBCs, highlighting the energy dependence of the process. DMSO alone had no effect on Gd uptake, whereas metabolic inhibition mimicked the effect of low temperature (blue). Data are shown as mean ± SD of three biological replicates. Statistical analysis was performed using one-way ANOVA followed by Bonferroni’s post hoc test. **P < 0.01.

### Gadoterate meglumine uptake is an endocytosis-independent process

The previously presented data strongly suggest that gadoterate meglumine uptake is an active, energy-dependent process. Among cellular mechanisms responsible for the internalization of extracellular molecules, endocytosis represents one of the most prominent pathways. To assess its involvement, K562 cells and WBCs were treated with the endocytosis inhibitors Dyngo 4a, Dynole 2-24, and chlorpromazine. Supplementary Figures S1-S3 show the range of concentrations tested to achieve effective endocytosis inhibition in K562 cells while maintaining low cytotoxicity. Inhibitory efficacy was evaluated using fluorescent transferrin and cholera toxin B subunit (CTxB). Transferrin uptake, which occurs via clathrin-mediated endocytosis, was strongly reduced by all three inhibitors (Dyngo 4a, Dynole 2-24, and chlorpromazine; Figs. S1–S3). In contrast, CTxB uptake, which occurs through lipid raft-mediated endocytosis, was inhibited by Dyngo 4a and Dynole 2-24 (Figs. S1, S2), but not by chlorpromazine (Fig. S3). Based on these assays, the optimal inhibitor concentrations for K562 cells under our experimental conditions were 50 µM Dyngo 4a, 30 µM Dynole 2-24, and 100 µM chlorpromazine.

Once the optimal concentration for each endocytosis inhibitor was established, K562 cells were co-incubated with the inhibitors and 2 mM gadoterate meglumine and analyzed by SC-ICP-MS. Figure 4A shows the intracellular Gd concentration after co-incubation with the different endocytosis inhibitors. Inhibition of endocytosis with Dynole 2-24, chlorpromazine, or Dyngo-4a did not reduce intracellular Gd levels after 1 h of incubation with 2 mM gadoterate meglumine.

**Figure 4.**
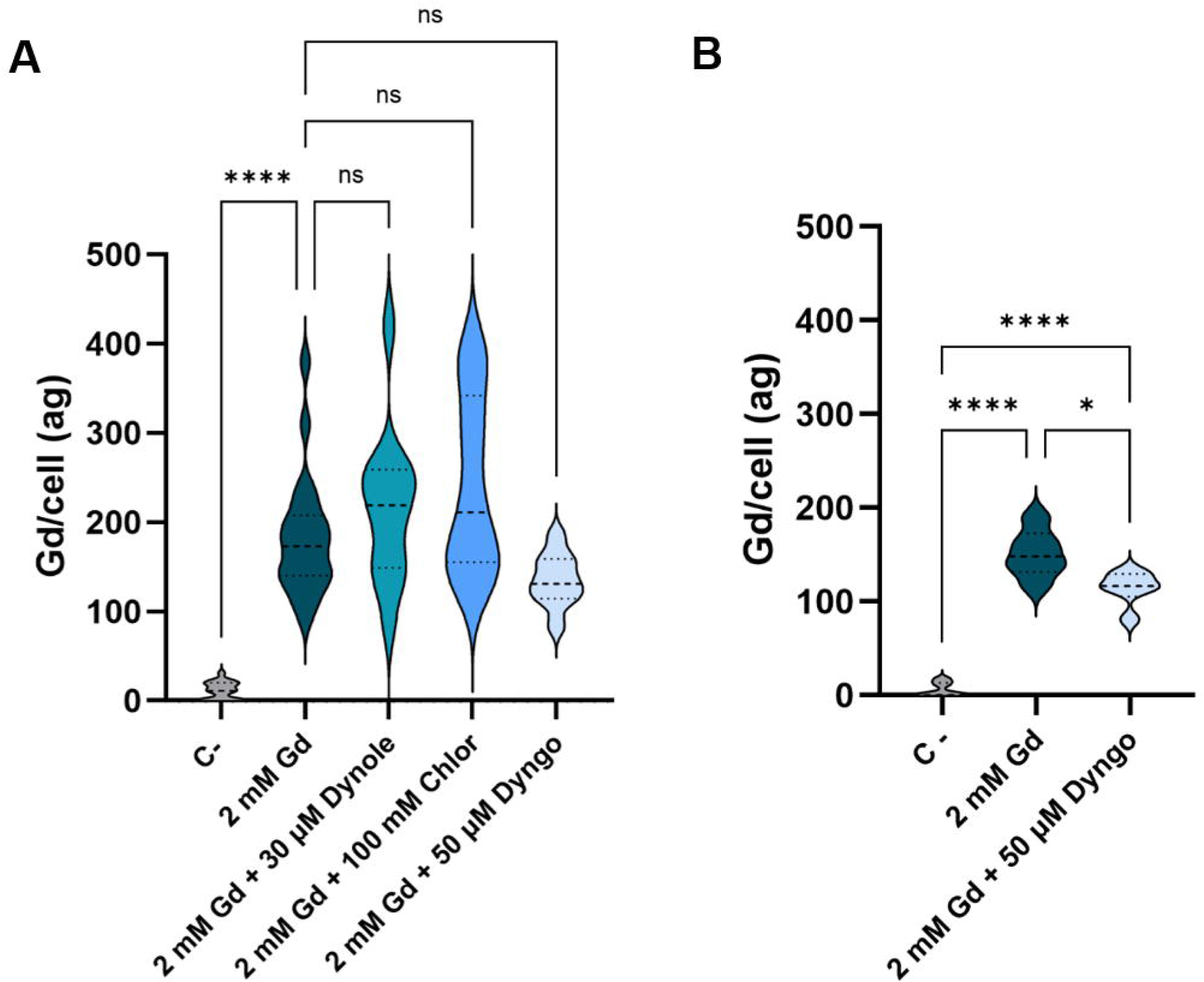
Endocytosis inhibition does not significantly alter gadoterate meglumine uptake in K562 cells (A) or WBCs (B). Uptake of gadoterate meglumine by K562 cells was not affected by inhibition of endocytosis with Dynole 2-24, chlorpromazine, or Dyngo-4a. In WBCs, treatment with Dyngo-4a caused a slight reduction in intracellular Gd levels; however, uptake remained substantially higher than in the negative control. Cells not incubated with gadoterate meglumine were used as negative controls (grey). Data are shown as mean ± SD of three biological replicates. Statistical analysis was performed using one-way ANOVA followed by Bonferroni’s post hoc test. *P < 0.1, ****P < 0.0001.

To further validate this result in WBCs, an additional experiment using Dyngo-4a was performed. Figure 4B shows a slight decrease in Gd uptake; however, this reduction is not biologically significant, indicating that endocytosis is unlikely to be the main entry mechanism of gadoterate meglumine in WBCs.

### Intracellular Retention and Active Release of Gadolinium in K562 Cells and WBCs

Our results show that, while decreasing its presence, Gd can be found intracellularly up to 24 hours after a 1 hour incubation with 2 mM gadoterate meglumine. In K562 cells the intracellular Gd concentration reduces by 83 ± 4% in 24h, following a one phase decay kinetic, while in human WBCs extracted from healthy donors, intracellular Gd concentration decreases by 73 ± 4% in 24h (Fig 5A,B; Table S1).

**Figure 5.**
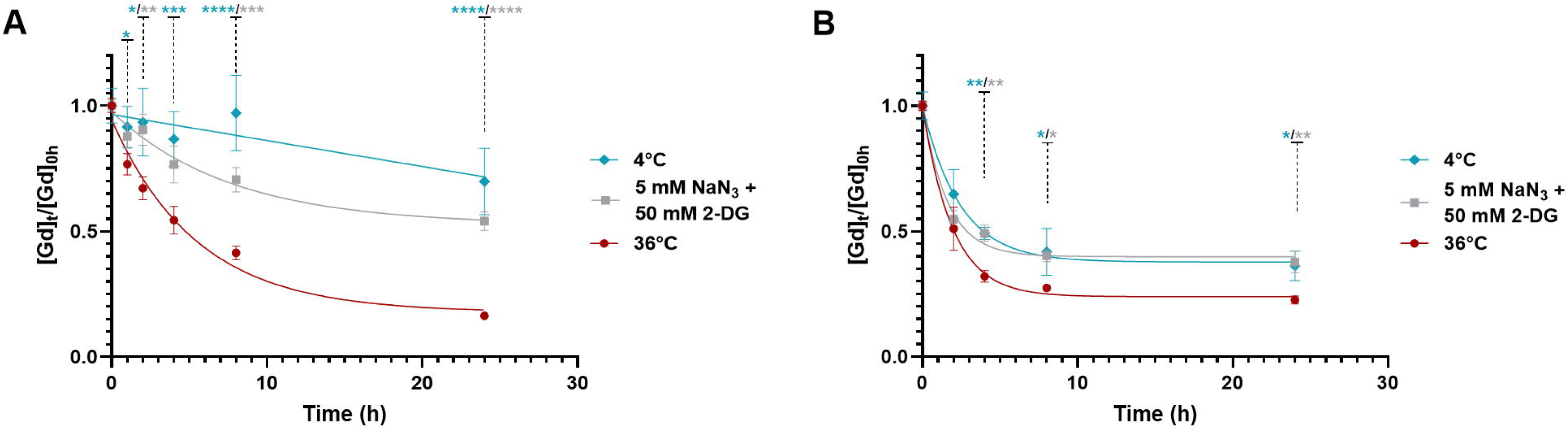
Gadolinium is retained intracellularly for up to 24 h and its release is an active process in K562 cells. **(A) and WBCs (B).** Under physiological conditions, intracellular Gd levels decrease following a one phase decay kinetic in both K562 and WBCs. In contrast, in K562 cells Gd release was strongly reduced under ATP-depleting conditions (5 mM NaN₃, 50 mM 2-deoxy-D-glucose) or when cellular metabolism was inhibited by low temperature (4 °C), highlighting the energy dependence of the process. Data are shown as mean ± SD of three biological replicates. **(B)** In WBCs, both treatments produced a mild decrease of the Gd release but less intense than K562 cells. Statistical analysis was performed using two-ways ANOVA followed by Bonferroni’s post hoc test. Mean values, SD and p values of the group comparisons are showed in Table S1. Blue asterisks indicate the p values for comparisons between the 4 °C condition and the control, whereas red asterisks indicate comparisons between the NaN₃ + 2-DG condition and the control. *P < 0.1, **P < 0.01, ***P < 0.001, ****P < 0.0001.

Furthermore, inhibition of cellular metabolism led to a significant reduction in Gd release. Figure 5A shows how both incubations with 5 mM NaN_3_ + 50 mM 2-DG and the K562 incubation at 4°C significantly reduced Gd externalization, with a more pronounced effect observed at 4 °C. The intracellular Gd concentration in metabolically inhibited K562 cells decreased by 30 ± 13 % at 4°C and by 46 ± 11% following incubation with NaN₃ plus 2-DG. Notably, K562 cells incubated at 4 °C exhibited a linear Gd release, deviating from the one-phase decay kinetics observed under control conditions. Similarly, inhibition of cellular metabolism in WBCs led to a reduced Gd release, with reductions of 64 ± 4% at 4°C and by 62 ± 10% in cells incubated with 5 mM NaN_3_ + 50 mM 2-DG (Fig 5A,B; Table S1). It is noteworthy that WBCs displayed a strong Gd release between 0 and 2 hours post-incubation, externalizing 40-50% of the total Gd within this short period, independently of the treatment applied. This early release may account for the comparatively milder effects observed under metabolic inhibition conditions.

Since extracellular vesicles (EVs) biogenesis is an active process and our results indicate that Gd release is also active, we assessed the presence of Gd in EVs shed by both K562 cells and WBCs. EV purification was successfully performed by ultracentrifugation followed by size-exclusion chromatography (SEC). Figure 6A,B shows the size distribution of the isolated EV population. EVs released by gadoterate meglumine-stimulated K562 cells showed a median size of 135 ± 13.8 nm while those shed by WBCs (Fig 6B) showed a median size of 154.7 ± 10.8 nm, comparable to non-stimulated K562 and WBCs, respectively (139.5 ± 9.3 nm 157.1 ± 6.1 nm). As a result of the SEC, 13 fractions were obtained. The EVs were eluted in fractions F3-F5, while the vast majority of the proteins were eluted in fractions F7-F11 (Fig 6C). The EV-containing fractions were pooled and analyzed by ICP-MS to assess whether EVs carried Gd. Fractions F1-F2 and F6-F13 were pooled and used as a control. Figure 6D shows the results of these measurements, indicating that EVs shed by WBCs were loaded with Gd 24 hours after cell incubation, whereas those shed by K562 cells were not (data not shown). Based on the Gd concentration measured by ICP-MS and the number of EVs determined by Nanoparticle Tracking Analysis (NTA), we estimate that EVs released by WBCs contain an average of 0.44 ag of Gd per EV. Notably, the supernatant obtained after ultracentrifugation was also analyzed and showed a Gd concentration of 2.9 µg L^-1^. This supernatant represents the non-particulate fraction of the WBC secretome, indicating that although Gd is released via EVs, this is not the only mechanism of transport to the extracellular milieu.

**Figure 6.**
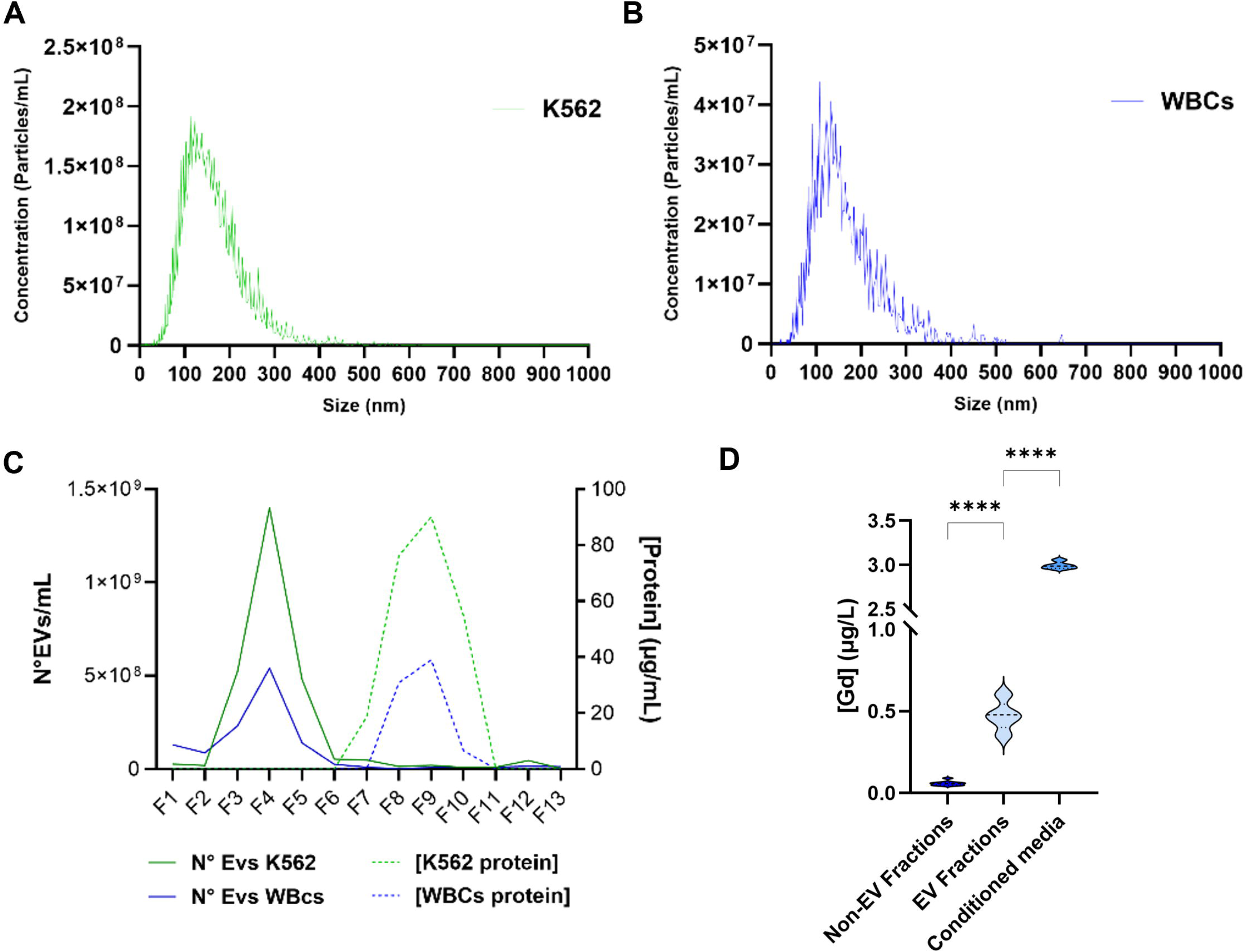
WBCs externalize gadolinium through their extracellular vesicles. Extracellular vesicles shed by K562 (A) and WBCs (B) were measured using NTA, showing median sizes of 135 ± 13.8 nm and 154.7 ± 10.8 nm, respectively. (C) Graph showing the EV and protein content in each of the 13 fractions obtained after SEC, indicating that EV-containing fractions have no contaminations from free protein. (D) ICP-MS measurements demonstrates that the EV fractions contain Gd, whereas non-EV fractions do not. Additionally, the conditioned media, consisting in the non-particulated secretome, showed a high concentration of Gd. Data are shown as mean ± SD of three technical replicates. Statistical analysis was performed using one-way ANOVA followed by Bonferroni’s post hoc test. ****P < 0.0001.

## Discussion

Administered intravenously, GBCAs first encounter cells of the vascular system and, in patients, is internalized by WBCs, as previously shown by our group (19). Once internalized within circulating cells, GBCAs may be distributed to virtually all tissues and organs, partially explaining the presence of Gd deposits in multiple organs observed in patients with NSF (10). Understanding the cellular uptake and retention of GBCAs by WBCs is an important area of investigation, given the ongoing concerns regarding long-term Gd deposition in human tissues (5, 25). A critical first step toward elucidating this process is determining whether GBCAs enter WBCs via passive or active transport mechanisms. Clarifying the nature of this uptake process is essential for understanding GBCA biodistribution, cellular retention, and potential downstream biological effects within immune cells (21). In this context, the present study investigated the cellular uptake mechanism of a clinically relevant concentration of gadoterate meglumine, a highly stable macrocyclic GBCA, using both primary WBCs and the human K562 leukemia cell line as experimental models.

To assess whether gadoterate meglumine uptake is mediated by a specific transporter, we first performed concentration-dependent uptake experiments in K562 cells and analyzed the data using a Michaelis-Menten kinetic model. This approach is well established for distinguishing between passive diffusion and carrier-mediated transport. Passive diffusion is characterized by a linear, non-saturable increase in uptake with increasing substrate concentration, whereas transporter-mediated uptake exhibits saturable kinetics, reaching a maximum velocity (Vmax) when transporters become fully occupied (26, 27). By fitting a concentration-dependent uptake data for gadoterate meglumine to the Michaelis-Menten model, we can calculate key kinetic parameters like the Michaelis constant (K_m_) and V_max_, providing definitive evidence for the involvement of a specific cellular transporter in the internalization of GBCAs. Our data revealed a clear saturable uptake profile for gadoterate meglumine in K562 cells, consistent with carrier-mediated transport. Fitting the data to the Michaelis–Menten equation yielded a V_max_ of 1.2×10^−11^ mol⋅g protein^−1^⋅s^−1^ and a K_m_ of 1.3 mM. This K_m_ value (low mM range) corresponds approximately to the value for the uptake of gadoxetic acid in hepatocytes by OATP1B1 or OATP1B3 (28).These findings were further supported by time-dependent uptake experiments performed at 2 mM gadoterate meglumine, a concentration commonly used in clinical practice, which showed a similar uptake behavior. In line with our results, De Aguero et al. (2025) reported similar kinetics in kidney cells incubated with 2 mM gadoterate meglumine, observing a plateau in intracellular accumulation at approximately 24 hours post-incubation (29). As unsynchronized cells were used in this study, we sought to assess whether uptake occurred preferentially in a specific phase of the cell cycle. However, our results indicate that drug internalization is independent of the cell cycle phase. Together, these results suggest that gadoterate meglumine internalization in K562 cells is mediated by a saturable transport mechanism rather than passive diffusion.

To further substantiate the involvement of active transport, we examined gadoterate meglumine uptake under energy-depleting conditions, including reduced temperature and metabolic inhibition. Incubation at sub-physiological temperatures resulted in a marked reduction in uptake at 16 °C and near-complete inhibition at 4 °C in both WBCs and K562 cells. Similarly, chemical inhibition of cellular metabolism using sodium azide and 2-deoxyglucose fully abolished gadoterate meglumine uptake. These findings provide direct evidence that gadoterate meglumine internalization is an energy-dependent process. Overall, the combined observations of time-, temperature-, and energy-dependent uptake, together with a saturable kinetic profile, collectively rule out simple passive diffusion as the primary entry mechanism.

Previous studies by Di Gregorio and colleagues proposed that GBCAs are internalized into fibroblasts and macrophages mainly via pinocytosis and subsequent endosomal sequestration (11). However, these conclusions were based on experiments performed using supraphysiological GBCA concentrations (5-100 mM) and prolonged exposure times (5-18 h), conditions that likely exaggerate vesicular uptake mechanisms compared to typical *in vivo* exposure scenarios. In contrast, our experiments were conducted using clinically relevant concentrations and incubation times. Importantly, co-incubation with endocytosis inhibitors, including Dyngo-4a and Dynole 2-24, which block dynamin-dependent vesicle scission, and chlorpromazine, which disrupts clathrin-mediated endocytosis, did not significantly reduce intracellular Gd levels in K562 cells and only marginally affected uptake in WBCs. These results show that endocytic pathways, including clathrin-mediated endocytosis, macropinocytosis, and phagocytosis, are not the primary routes of gadoterate meglumine entry into these cells.

Collectively, our findings support the existence of an alternative, non-endocytic transport pathway and strongly suggest that gadoterate meglumine uptake in both K562 cells and WBCs is mediated by an as-yet unidentified membrane transporter. This conclusion aligns with growing evidence for other GBCAs. Notably, uptake of the hepatobiliary GBCA gadoxetate into hepatocytes has been shown to be saturable and mediated specifically by the organic anion-transporting polypeptide OATP1 (30). In that case, uptake could be competitively inhibited by known OATP substrates such as bromosulfophthalein and taurocholate, providing clear evidence for carrier-mediated transport (31). Although gadoterate meglumine and gadoxetate differ in chemical structure and biodistribution, these findings demonstrate that GBCAs, despite their hydrophilic nature, can be actively transported into cells via specific membrane proteins. This further supports our conclusion that gadoterate meglumine uptake is an active and regulated cellular process.

As mentioned, previous work from our group demonstrated that Gd can be detected in patients following gadoterate meglumine–enhanced MRI and that WBCs from some patients accumulate significantly higher GBCA levels than others (19). We have also shown that circulating immune cells can internalize GBCAs without impairing their ability to migrate across the blood–brain barrier *in vitro* (20). These observations suggest that immune cells may act as vehicles for Gd distribution and tissue deposition. To shed light into the downstream consequences of the Gd uptake, we assessed the Gd release by K562 cells and WBCs. To this end, cells were incubated with 2 mM gadoterate meglumine for 1 hour at 36°C, washed to remove non-internalized compound, and intracellular Gd levels were subsequently measured. In both cell types, a substantial fraction of the internalized Gd was released over a 24-hour period. Moreover, our results demonstrate that Gd release is predominantly mediated by an active process in K562 cells and, at least partially, by active mechanisms in WBCs.

The chemical form in which Gd is externalized remains unclear. In particular, whether it is released as intact gadoterate meglumine or undergoes transmetalation or transligandation, respectively, represents a key open question in the field (11, 32). This distinction is likely to determine the capacity of Gd to be re-internalized by other cell types, thereby contributing to its long-term accumulation in tissues. Notably, a small fraction of Gd remains intracellular even 24 hours after stimulation in both K562 cells and WBCs. Intracellular Gd has been reported to persist in other cell types in the form of insoluble precipitates, the precise composition of which remains unresolved; however, co-localization with phosphorus and oxygen has led to the hypothesis that GdPO₄-like deposits may form (33). Importantly, in this work we demonstrate for the first time that WBCs can release Gd via extracellular vesicles, a mechanism that may prolong Gd circulation time in the bloodstream and enhance its potential to reach distal and immunoprivileged tissues (34). These findings suggest that Gd-loaded EVs could represent a mechanism for Gd entry into brain tissue; accordingly, further experiments are currently underway in our laboratory to investigate this hypothesis. In line with this, *in vitro*-engineered EVs have been shown to mediate the transfer of Gd to different tissues and cell types, partly escaping from renal clearance due to their biomimetic properties and nanoscale size (∼130 nm), which limits glomerular filtration (35). Furthermore, the capacity of EVs to mediate intercellular metal transfer has recently been demonstrated by Purnianto et al., who reported the transport of both toxic and non-toxic metals between distinct cell populations, therefore, the potential of these EVs to transfer Gd cannot be overlooked (36). Moreover, EVs have been described to play important roles in both bone homeostasis and bone-related pathologies (37, 38). The potential impact of Gd-containing EVs shed by WBCs, osteoblasts, and osteoclasts on bone formation and resorption remains an open question in the field, which our laboratory is beginning to address.

Building on these findings, the present study provides the first mechanistic framework by demonstrating that gadoterate meglumine uptake and release are mediated by active processes. Importantly, this implies that GBCA internalization and secretion are regulated and may vary depending on transporter expression, cellular metabolic state, or immune activation status. Such variability could contribute to the heterogeneous GBCA loads observed in patients, particularly in immunocompromised individuals or those with gadolinium deposition disease (GDD). In patients with chronic inflammatory conditions or altered immune cell function, changes in immune cell metabolism, trafficking, or transporter expression could directly influence GBCA uptake and redistribution, potentially increasing long-term Gd retention (39). Furthermore, understanding the role of Gd-loaded extracellular vesicles in its dissemination across different tissues, including immunoprivileged sites, would represent a major advance in elucidating the pathophysiology of GDD. Finally, identifying the transporter(s) responsible for gadoterate meglumine internalization therefore represents a critical next step. Such knowledge would enable the rational screening of transport inhibitors and could open a new avenue for GBCA optimization, including the co-administration of uptake blockers to minimize cellular internalization and reduce long-term gadolinium retention in patients. Altogether, these findings reinforce the concept that GBCAs are not inert agents, but dynamically handled substrates of cellular physiology, with potential implications for their safety, biodistribution, and long-term fate in the human body.

## Materials and methods

### Cell culture

For all cell-based experiments, the human myelogenous leukemia cell line K562 (ATCC Catalog #CCL-243™) was utilized as a model due to its origin from blood-forming tissue and non-adherent nature. K562 cells were cultured in RPMI 1640 w/o Phenol Red and w/o riboflavin, supplemented with 2 mM stabilized L-Glutamine, 4.5 mM HEPES, (BioConcept, Allschwil, Switzerland) and 10% FBS (Gibco, Grand Island, NY, USA), and maintained in a humidified incubator with 3.5% CO_2_ at 36°C.

### Blood collection and white blood cell isolation

For this study, blood samples were collected from healthy, consented donors into heparinized collection tubes. WBCs were isolated via erythrocyte lysis using eBioscience™ 10X RBC Lysis Buffer (Multi-species) (10X), (Invitrogen, Waltham, MA, USA) solution. Cells were then washed two times with RPMI 1640 and finally resuspended in RPMI 1640 for further analysis.

### GBCA loading studies

Clinically, patients are administered GBCAs intravenously at a concentration of 2mM for contrast enhanced MRIs which is then eliminated by the kidneys with an early elimination half-life of less than 2 hours in patients with normal renal function, >95% of the GBCA is cleared from the body in 12 hours (40). Commercialized form of Gd-DOTA Dotarem - Gadoterate Meglumine, (Guerbet, Villepinte, France) was obtained from the Insel Hospital Pharmacy (Bern, Switzerland).

- Concentration-dependent uptake

For the kinetic analysis of gadoterate meglumine uptake by K562 cells, 500 µL of a cell suspension (1 × 10⁷ cells/mL) was incubated with increasing concentrations of gadoterate meglumine (0.2–10 mM) for 60 minutes at 36 °C. Cells were centrifuged at 300 × g for 3 minutes and washed with PBS supplemented with 0.5% BSA. Cells were then fixed with 1% PFA for 20 minutes with gentle mixing by inversion. Fixed cells were subsequently centrifuged and washed three times with PBS + 0.5% BSA, and finally resuspended in PBS w/o calcium or magnesium ions. Intracellular Gd content was quantified by SC-ICP-MS (see below).

- Time-dependent uptake

A 10 mL cell culture (1 × 10⁶ cells/mL) was stimulated with 2 mM gadoterate meglumine and incubated in a humidified atmosphere containing 3.5% CO₂ at 36 °C. At each experimental time point (0, 1, 2, and 6 h), 1 mL of the cell suspension was collected. Cells were then washed and fixed as previously described. Intracellular gadolinium (Gd) content was subsequently quantified by single-cell inductively coupled plasma mass spectrometry (SC-ICP-MS) (see below).

- Temperature-dependent uptake

The analysis of the influence of temperature on Gd uptake was performed in a similar manner. Both K562 cells and WBCs were incubated with 2 mM gadoterate meglumine for 1 hour. Three different temperatures were tested: 36 °C (in an incubator, serving as the positive control), 16 °C (in an incubator), and 4 °C (on ice). Cells were then washed and fixed as previously described. Gd content was quantified by SC-ICP-MS (see below).

- Energy-dependent uptake

Cellular energy production was inhibited by pre-treating the cells for 30 min with 5 mM NaN_3_ (Sigma Aldrich, St. Louis, MO, USA) to inhibit oxidative phosphorylation and 50 mM 2-deoxy-D-glucose (Sigma Aldrich, St. Louis, MO, USA) to block glycolysis. Subsequently, 2 mM gadoterate meglumine was added and the incubation continued for 1 hour. Then the cells were fixed with paraformaldehyde, washed with PBS and processed for the measurement of Gd by SC-ICP-MS.

- Endocytosis-mediated uptake

The potential involvement of endocytosis in Gd uptake was evaluated by pharmacological inhibition using Dyngo 4a (Monmouth Junction, NJ, USA), Dynole 2-24 (Monmouth Junction, NJ, USA), and chlorpromazine (Sigma Aldrich, St. Louis, MO, USA). The optimal inhibitor concentrations required to achieve effective blockade of endocytic pathways were determined by fluorescence microscopy through co-incubation with Alexa Fluor 555-transferrin (Invitrogen, Waltham, MA, USA) and Alexa Fluor 555-cholera toxin B subunit (CTB) (Invitrogen, Waltham, MA, USA). Briefly, 1 × 10⁵ K562 cells were seeded onto sterilized coverslips pre-treated with poly-D-lysine (Gibco, Grand Island, NY, USA) and incubated overnight under standard culture conditions. Cells were then washed three times with pre-warmed RPMI medium w/o FBS to eliminate residual transferrin present in the serum. Subsequently, cells were pre-incubated with increasing concentrations of endocytosis inhibitors (0.5-100 µM) for 30 minutes, followed by the addition of 25 µg/mL Alexa Fluor 555–transferrin or 0.6 µg/mL Alexa Fluor 555-CTB for an additional 30 minutes. Cells were then washed three times with PBS supplemented with calcium and magnesium and fixed with 0.5% paraformaldehyde for 30 minutes. Images were acquired using an inverted EVOS™ M5000 Imaging System (Invitrogen, Waltham, MA, USA). The same methodology and experimental conditions were applied for the analysis of WBCs.

Once the optimal inhibitor concentrations were selected to maximize endocytosis inhibition while minimizing cellular stress, K562 cells were co-incubated for 1 hour with 2 mM gadoterate meglumine in the presence of either 50 µM Dyngo 4a, 30 µM Dynole 2-24, or 100 µM chlorpromazine. Cells treated with 2 mM gadoterate meglumine alone, with or without DMSO, were used as positive controls, whereas cells incubated without gadoterate meglumine served as negative controls. All procedures were performed following the methodology described above. The same methodology and experimental conditions were applied for the analysis of WBCs. Gd content was quantified by SC-ICP-MS (see below).

### Cell cycle analysis

To analyze the influence of the cell cycle phase on gadoterate meglumine uptake, 10^7^ unsynchronized cells were incubated with 2 mM gadoterate meglumine at 36°C for 1 hour, as described above. After incubation, cells were washed three times and stained with 5 µg/mL DRAQ5 (Thermo Fisher, Waltham, MA, USA) for 10 minutes at 36°C in the dark. Subsequently, cells were sorted based on their DNA content into G0/G1, S, and G2/M phases using a BD FACSDiscover S8 cell sorter (BD Biosciences, Franklin Lakes, NJ, USA). Sorted cells were fixed with 1% PFA for 20 minutes, washed 3 times with PBS + 0.5% BSA and resuspended in PBS. Intracellular Gd content was quantified by SC-ICP-MS.

### Gadolinium release analysis

To determine the intracellular retention time of Gd and to assess if its externalization is an active process, 3 x 10^7^ K562 cells were stimulated with 2 mM gadoterate meglumine and subsequently washed as previously described. Once cells were washed, they were divided into three experimental conditions: (i) 10^7^ cells were resupended into RPMI + 10% FBS and incubated at 36°C; (ii) 10^7^ cells were resuspended in RPMI + 10% FBS + 5 mM sodium azide and 50 mM 2-Deoxyglucose and incubated at 36°C; (iii) 10^7^ cells were resuspended in RPMI + 10% FBS and incubated at 4°C. At the indicated time points (0, 1, 2, 4, 8, and 24 h), 10^6^ cells were collected from each condition and fixed as previously described. Intracellular Gd content was quantified by SC-ICP-MS.

### Extracellular vesicles purification

To assess whether Gd is released via extracellular vesicles (EVs), conditioned media were collected 24 hours after the gadoterate meglumine incubation. The conditioned media were centrifuged at 1500 xg for 15 minutes at 4°C to remove cell debris. The supernatant was then ultracentrifuged at 120000 xg for 3 hours at 4°C using the TH-641 Swinging Bucket Rotor in a Sorvall 100WX+ ultracentrifuge (Thermo Fisher, Waltham, MA, USA). The EV pellet was resuspended in 1 mL of 0.22 µm-filtered PBS. The EVs suspension was then subjected to a size exclusion chromatography using a 20 mL sepharose column (Izon Science, Christchurch, Canterbury, New Zealand). The collected fractions were analyzed by Nanoparticle Tracking Analysis using a Zetasizer instrument (Malvern Panalytical, Worcestershire, UK). EVs were lysed by sonication and the Gd content in both the EV lysates and the conditioned media supernatant was quantified by conventional ICP-MS.

### Single-Cell Inductively Coupled Plasma Mass Spectrometry (SC-ICP-MS)

All single-cell inductively coupled plasma mass spectrometry (SC-ICP-MS) analyses were performed using a NexION 2000 ICP-MS (PerkinElmer, Shelton, CT, USA) as previously described (41). The instrument was equipped with an Asperon spray chamber (PerkinElmer Catalog #N8152493), a 100 µm nebulizer, and an integrated autosampler for automated sample introduction. The ICP-MS was operated in single-cell acquisition mode, optimized for the detection of transient signals generated by individual particles or cells. Argon gas was used as the plasma gas at a flow rate of 15-17 L/min and as the nebulizer gas at a flow rate of 0.4 L/min. The dwell time per isotope was set to 50 µs, with an acquisition time of 120 seconds per sample.

Instrument Performance and Calibration - Prior to each experiment, an instrument performance check was conducted. Multi-element calibration standards - NexION Dual Detector Solution, 2% HNO3 (PerkinElmer, Shelton, CT, USA) and NexION Setup Solution, 1 μg/L (PerkinElmer, Shelton, CT, USA) containing beryllium (Be), indium (In), and uranium (U) were analyzed to verify the instrument’s detection capabilities across a wide range of atomic masses. This ensured optimal sensitivity and long-term stability for accurate quantitative analysis. The plasma health was further assessed by monitoring critical oxide and double-charge formation ratios (CeO/Ce and Ce^2+^/Ce). These ratios are key indicators of plasma conditions and potential spectral interferences. Transport efficiency was consistently determined using freshly prepared 50 nm Ultra Uniform Gold Nanospheres (PerkinElmer, Shelton, CT, USA). Immediately prior to analysis, the gold nanoparticle suspension was sonicated for 10 minutes at 20 °C to ensure uniform dispersion. This measurement was performed at a sample flow rate of 0.01 mL/min with a particle concentration of 110,000 particles/mL, allowing for accurate quantification of nebulization and transport efficiency.

The SC-ICP-MS was calibrated for the quantitative analysis of ^24^Mg and ^157^Gd. Calibration standards with defined concentrations of 1, 5, 10, and 20 ppb were prepared in 2% HNO_3_ A calibration curve was generated for each isotope, and experiments were only initiated when the coefficient of determination (*R^2^*) for these curves exceeded 0.9, ensuring the linearity and reliability of the calibration.

Sample Introduction and Isotope Monitoring - Cell suspensions were introduced directly into the nebulizer by aspiration at a controlled sample flow rate of approximately 0.05 mL/min. To optimize signal acquisition, samples were diluted to a target cell concentration of approximately 10^5^ cells/mL. For gadolinium detection, the ^157^Gd isotope was specifically monitored. Further, ^24^Mg was monitored across all experiments, serving as an internal cellular presence control to confirm successful introduction and detection of individual cells. Referencing the work of Di Gregorio and colleagues, who demonstrated that macrocyclic GBCAs remain intact in WBCs for up to 3 hours, we assume the GBCAs in our analysis are part of the chelate. Consequently, all elemental gadolinium measures subsequently are denoted as Gd (20, 42).

### Statistical analysis

All the experiments included at least three biological replicates, each of them consisting in three technical replicates. All quantitative data are presented as the mean ± standard deviation (SD). For comparisons involving multiple groups, a one-way analysis of variance (ANOVA) with repeated measures was performed. Specific statistical tests used and corresponding *p*-values for individual comparisons are reported in the respective figure legends. All statistical analyses were conducted using GraphPad Prism software version 10.4.1 (GraphPad Software, La Jolla, CA, USA).

## Acknowledgments

The authors would like to thank the radiology technicians of the Radiology Department at Inselspital for performing the blood collection. We also acknowledge the technical support and assistance with the laboratory experiments provided by Nicola Lüdi and the Flow Cytometry and Cell Sorting Facility (FCCS) of the DBMR (University of Bern). Finally, the authors would also like to acknowledge the Master’s students Ilia Jenny and Sophia Cestaroni, as well as the Bachelor’s students Gaia Bignasca and Talin Aebi, for their support with laboratory tasks during this period.

## Supporting Information

**Supplementary Figure S 1 Endocytosis inhibition in K562 cells by Dyngo 4a.** Cells were incubated with Dyngo 4a for 1 hour, and endocytic activity was assessed using Alexa Fluor 555-transferrin (A) and Alexa Fluor 555-CTxB (B). Dyngo 4a efficiently reduced the uptake of both transferrin and CTxB, with the strongest inhibition observed at 50 µM. Therefore, 50 µM was selected as the optimal concentration for subsequent experiments.

**Supplementary Figure S 2 Endocytosis inhibition in K562 cells by Dynole 2-24.** Cells were incubated with Dynole 2-24 for 1 hour, and endocytic activity was assessed using Alexa Fluor 555-transferrin (A) and Alexa Fluor 555-CTxB (B). Dynole 2-24 efficiently reduced the uptake of both transferrin and CTxB, with the strongest inhibition observed at 50 µM; however, this concentration also decreased cell viability. Therefore, 30 µM was selected as the optimal concentration for subsequent experiments. Notably, a subset of cells displayed very high CTxB binding, and these extremely-positive cells were not affected by Dynole 2-24 treatment, likely corresponding to apoptotic or necrotic cells.

**Supplementary Figure S 3 Endocytosis inhibition in K562 cells by chlorpromazine.** Cells were incubated with chlorpromazine for 1 hour, and endocytic activity was assessed using Alexa Fluor 555-transferrin (A) and Alexa Fluor 555-CTxB (B). Chlorpromazine efficiently reduced the uptake of transferrin and reduces the uptake of CTxB, with the strongest reduction observed at 100 µM. Again, a subset of cells displayed very high CTxB binding which were unaffected by the treatment, likely corresponding to apoptotic or necrotic cells.

**Supplementary Table 1 Statistical analysis was performed using two-ways ANOVA followed by Bonferroni’s post hoc test. Data are shown as mean ± SD of three biological replicates.**

---

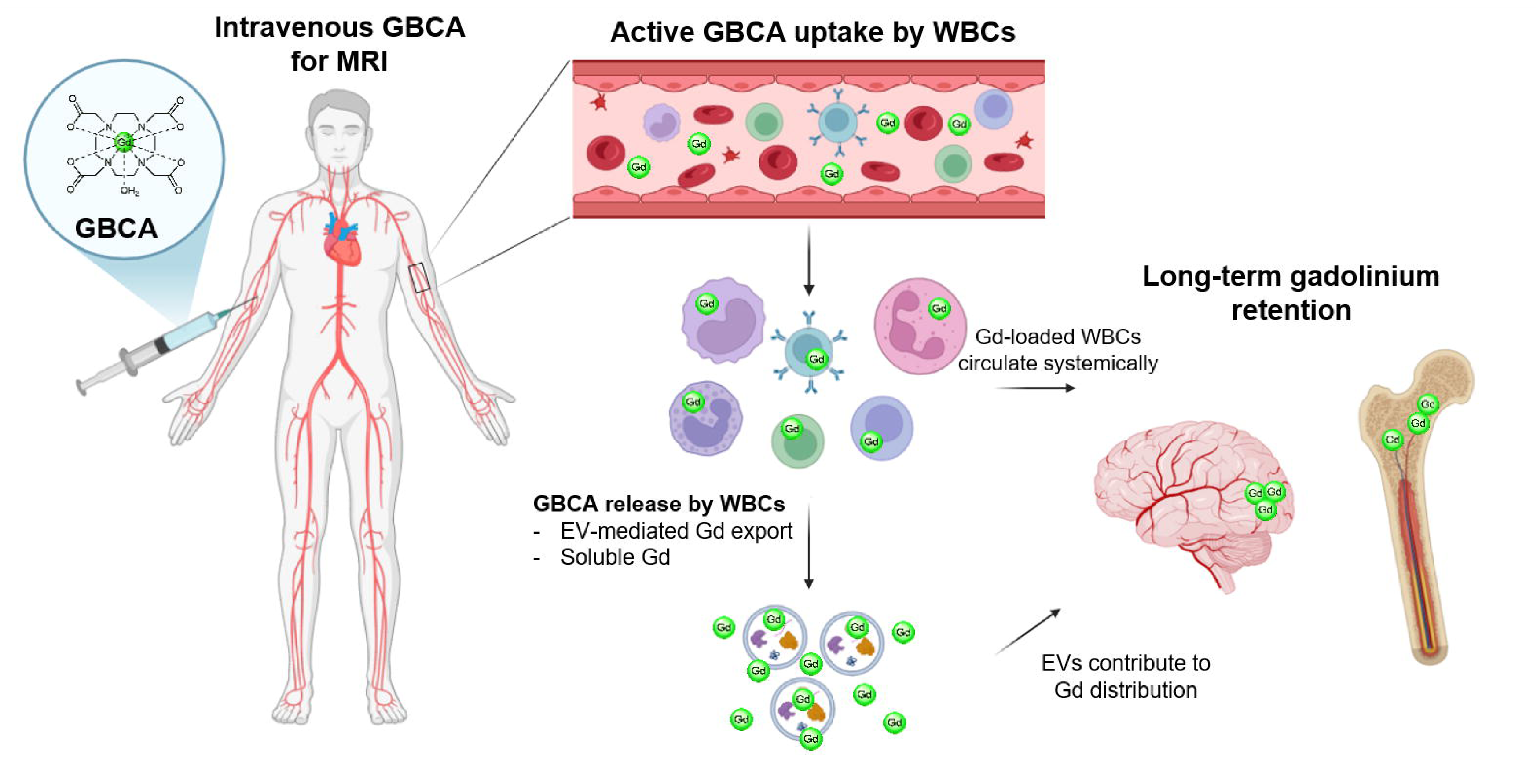

## References

1. Lauterbur PC. Image formation by induced local interactions. Examples employing nuclear magnetic resonance. 1973. Clin Orthop Relat Res. 1989(244):3–6.

2. Yousaf T, Dervenoulas G, Politis M. Advances in MRI Methodology. Int Rev Neurobiol. 2018;141:31–76.

3. Dekker HM, Stroomberg GJ, Van der Molen AJ, Prokop M. Review of strategies to reduce the contamination of the water environment by gadolinium-based contrast agents. Insights Imaging. 2024;15(1):62.

4. Weinreb JC, Rodby RA, Yee J, Wang CL, Fine D, McDonald RJ, et al. Use of Intravenous Gadolinium-Based Contrast Media in Patients With Kidney Disease: Consensus Statements from the American College of Radiology and the National Kidney Foundation. Kidney Med. 2021;3(1):142–50.

5. Domingo JL, Semelka RC. Gadolinium toxicity: mechanisms, clinical manifestations, and nanoparticle role. Arch Toxicol. 2025;99(10):3897–916.

6. Davies J, Siebenhandl-Wolff P, Tranquart F, Jones P, Evans P. Gadolinium: pharmacokinetics and toxicity in humans and laboratory animals following contrast agent administration. Arch Toxicol. 2022;96(2):403–29.

7. Behzadi AH, Zhao Y, Farooq Z, Prince MR. Immediate Allergic Reactions to Gadolinium-based Contrast Agents: A Systematic Review and Meta-Analysis. Radiology. 2018;286(2):471–82.

8. Cowper SE, Robin HS, Steinberg SM, Su LD, Gupta S, LeBoit PE. Scleromyxoedema-like cutaneous diseases in renal-dialysis patients. Lancet. 2000;356(9234):1000–1.

9. Grobner T. Gadolinium--a specific trigger for the development of nephrogenic fibrosing dermopathy and nephrogenic systemic fibrosis? Nephrol Dial Transplant. 2006;21(4):1104–8.

10. Sanyal S, Marckmann P, Scherer S, Abraham JL. Multiorgan gadolinium (Gd) deposition and fibrosis in a patient with nephrogenic systemic fibrosis--an autopsy-based review. Nephrol Dial Transplant. 2011;26(11):3616–26.

11. Di Gregorio E, Gianolio E, Stefania R, Barutello G, Digilio G, Aime S. On the fate of MRI Gd-based contrast agents in cells. Evidence for extensive degradation of linear complexes upon endosomal internalization. Anal Chem. 2013;85(12):5627–31.

12. Robert P, Violas X, Grand S, Lehericy S, Idee JM, Ballet S, et al. Linear Gadolinium-Based Contrast Agents Are Associated With Brain Gadolinium Retention in Healthy Rats. Invest Radiol. 2016;51(2):73–82.

13. Robert P, Fingerhut S, Factor C, Vives V, Letien J, Sperling M, et al. One-year Retention of Gadolinium in the Brain: Comparison of Gadodiamide and Gadoterate Meglumine in a Rodent Model. Radiology. 2018;288(2):424–33.

14. Gulani V, Calamante F, Shellock FG, Kanal E, Reeder SB, International Society for Magnetic Resonance in M. Gadolinium deposition in the brain: summary of evidence and recommendations. Lancet Neurol. 2017;16(7):564–70.

15. Ramalho J, Ramalho M, Semelka RC. Gadolinium Elimination in a Gadolinium Deposition Disease Population After a Single Exposure to Gadolinium-Based Contrast Agents. Invest Radiol. 2025;60(7):429–33.

16. McDonald RJ, Weinreb JC, Davenport MS. Symptoms Associated with Gadolinium Exposure (SAGE): A Suggested Term. Radiology. 2022;302(2):270–3.

17. Ponrartana S, Moore MM, Chan SS, Victoria T, Dillman JR, Chavhan GB. Safety issues related to intravenous contrast agent use in magnetic resonance imaging. Pediatr Radiol. 2021;51(5):736–47.

18. Lancelot E, Raynaud JS, Desche P. Current and Future MR Contrast Agents: Seeking a Better Chemical Stability and Relaxivity for Optimal Safety and Efficacy. Invest Radiol. 2020;55(9):578–88.

19. Ruprecht N, Parakkattel D, Hofmann L, Broekmann P, Ludi N, Kempf C, et al. Uptake of Gadolinium-Based Contrast Agents by Blood Cells During Contrast-Enhanced MRI Examination. Invest Radiol. 2024;59(5):372–8.

20. Parakkattel D, Ruprecht N, Broekmann P, Guimbal S, Studle C, Soldati S, et al. Identifying a potential role of immune cells in gadolinium deposition within the brain. Fluids Barriers CNS. 2025;22(1):80.

21. Roberts AG. The Structure and Mechanism of Drug Transporters. Methods Mol Biol. 2021;2342:193–234.

22. Stillwell W. Membrane Transport. An Introduction to Biological Membranes (Second Edition): Elsevier; 2016. p. 423–51.

23. Kierans SJ, Taylor CT. Glycolysis: A multifaceted metabolic pathway and signaling hub. J Biol Chem. 2024;300(11):107906.

24. Mader A. Glycolysis and oxidative phosphorylation as a function of cytosolic phosphorylation state and power output of the muscle cell. Eur J Appl Physiol. 2003;88(4-5):317–38.

25. Fretellier N, Idee JM, Rasschaert M, Factor C, Van der Molen AJ. Gadolinium Deposition in Bone Tissues After Contrast-enhanced Magnetic Resonance Imaging: A Comprehensive Review. Invest Radiol. 2025;60(11):722–44.

26. Michaelis L, Menten ML, Johnson KA, Goody RS. The original Michaelis constant: translation of the 1913 Michaelis-Menten paper. Biochemistry. 2011;50(39):8264–9.

27. Schnell S. Validity of the Michaelis-Menten equation--steady-state or reactant stationary assumption: that is the question. FEBS J. 2014;281(2):464–72.

28. Leonhardt M, Keiser M, Oswald S, Kuhn J, Jia J, Grube M, et al. Hepatic uptake of the magnetic resonance imaging contrast agent Gd-EOB-DTPA: role of human organic anion transporters. Drug Metab Dispos. 2010;38(7):1024–8.

29. DeAguero J, Howard T, Escobar GP, Dokladny K, Wagner B. Early endolysosomal dysfunction is a contributing factor to gadolinium-based contrast agent mouse renal proximal tubule epithelial cell injury. Cell Biol Toxicol. 2025;41(1):65.

30. Nassif A, Jia J, Keiser M, Oswald S, Modess C, Nagel S, et al. Visualization of hepatic uptake transporter function in healthy subjects by using gadoxetic acid-enhanced MR imaging. Radiology. 2012;264(3):741–50.

31. van Montfoort JE, Stieger B, Meijer DK, Weinmann HJ, Meier PJ, Fattinger KE. Hepatic uptake of the magnetic resonance imaging contrast agent gadoxetate by the organic anion transporting polypeptide Oatp1. J Pharmacol Exp Ther. 1999;290(1):153–7.

32. Frenzel T, Lengsfeld P, Schirmer H, Hutter J, Weinmann HJ. Stability of gadolinium-based magnetic resonance imaging contrast agents in human serum at 37 degrees C. Invest Radiol. 2008;43(12):817–28.

33. DeAguero J, Howard T, Kusewitt D, Brearley A, Ali AM, Degnan JH, et al. The onset of rare earth metallosis begins with renal gadolinium-rich nanoparticles from magnetic resonance imaging contrast agent exposure. Sci Rep. 2023;13(1):2025.

34. Yanez-Mo M, Siljander PR, Andreu Z, Zavec AB, Borras FE, Buzas EI, et al. Biological properties of extracellular vesicles and their physiological functions. J Extracell Vesicles. 2015;4:27066.

35. Rayamajhi S, Marasini R, Nguyen TDT, Plattner BL, Biller D, Aryal S. Strategic reconstruction of macrophage-derived extracellular vesicles as a magnetic resonance imaging contrast agent. Biomater Sci. 2020;8(10):2887–904.

36. Purnianto A, Mawal C, Kulkarni MM, Su H, Koukoulis TF, Wongsodirdjo P, et al. Small extracellular vesicles contain metals and transfer metal intercellularly. J Extracell Biol. 2024;3(10):e70012.

37. Liu M, Sun Y, Zhang Q. Emerging Role of Extracellular Vesicles in Bone Remodeling. J Dent Res. 2018;97(8):859–68.

38. Fang F, Yang J, Wang J, Li T, Wang E, Zhang D, et al. The role and applications of extracellular vesicles in osteoporosis. Bone Res. 2024;12(1):4.

39. Wiginton CD, Kelly B, Oto A, Jesse M, Aristimuno P, Ernst R, et al. Gadolinium-based contrast exposure, nephrogenic systemic fibrosis, and gadolinium detection in tissue. AJR Am J Roentgenol. 2008;190(4):1060–8.

40. van der Molen AJ, Quattrocchi CC, Mallio CA, Dekkers IA, European Society of Magnetic Resonance in Medicine BGREC. Ten years of gadolinium retention and deposition: ESMRMB-GREC looks backward and forward. Eur Radiol. 2024;34(1):600–11.

41. Gale A, Hofmann L, Ludi N, Hungerbuhler MN, Kempf C, Heverhagen JT, et al. Beyond Single-Cell Analysis of Metallodrugs by ICP-MS: Targeting Cellular Substructures. Int J Mol Sci. 2021;22(17).

42. Di Gregorio E, Furlan C, Atlante S, Stefania R, Gianolio E, Aime S. Gadolinium Retention in Erythrocytes and Leukocytes From Human and Murine Blood Upon Treatment With Gadolinium-Based Contrast Agents for Magnetic Resonance Imaging. Invest Radiol. 2020;55(1):30–7.

